# Geographic distribution and future expansion of *Aedes albopictus* in the Democratic Republic of the Congo

**DOI:** 10.1101/2021.10.22.465397

**Authors:** Fabien Vulu, Thierry Lengu Bobanga, Toshihiko Sunahara, Kyoko Futami, Hu Jinping, Noboru Minakawa

**Affiliations:** Program for Nurturing Global Leaders in Tropical and Emerging Communicable Diseases, Graduate School of Biomedical Sciences, Nagasaki University, Nagasaki, Japan; Vector Ecology & Environment Department, Institute of Tropical Medicine, Nagasaki University, Nagasaki, Japan; Services de Parasitologie et d’Entomologie, Département de Médecine Tropicale, Faculté de Médecine, Université de Kinshasa, Democratic Republic of the Congo

**Author notes:** Corresponding author: Fabien Vulu.

**Keywords:** *Aedes* mosquito, maximum entropy model, MaxEnt, environmental variables

## Abstract

*Aedes albopictus* with an Asian origin has been reported from central African countries. The establishment of this mosquito species poses a serious threat as the vector of various infectious diseases. Since information about *Ae. albopictus* in Democratic Republic of the Congo (DRC) is scarce, we investigated the current distribution of this mosquito species. Based on the factors affecting the distribution, we predicted future distribution. We conduced entomological surveys in Kinshasa and three neighboring cities from May 2017 to September 2019. The survey was extended to seven inland cities. A total of 19 environmental variables were examined using the maximum entropy method to identify areas suitable for *Ae. albopictus* to establish a population. We found *Ae. albopictus* at 21 of 23 sites in Kinshasa and three neighboring cities. For the first time *Ae. albopictus* was also found from three of seven inland cities, while it was not found in four cities located in the eastern and southeastern parts of DRC. A maximum entropy model revealed that the occurrence of *Ae. albopictus* was positively associated with maximum temperature of the warmest month, and negatively associated with wider mean diurnal temperature range and enhanced vegetation index. The model predicted that most parts of DRC are suitable for the establishment of the mosquito. The unsuitable areas were the eastern and southeastern highlands, which have low temperatures and long dry seasons. We confirmed that *Ae. albopictus* is well established in Kinshasa and its neighboring cities. The expansion of *Ae. albopictus* to the inland is ongoing, and in the future the mosquito may establish in most parts of DRC.

## Introduction

*Aedes albopictus* is an invasive mosquito and vector of human disease such arboviruses such as dengue and chikungunya arboviruses [1-5]. Originating from Asia [6, 7], *Ae. albopictus* has expanded its distribution globally [3]. In central Africa, this mosquito was first reported from Cameroon in 2000 [8], and subsequently was found in several other countries [9-13]. Following the mosquito invasion into central Africa, numerous dengue and chikungunya outbreaks have occurred [12, 14-21].

*Aedes aegypti* is considered to be the main vector of dengue and chikungunya (CHIKV) viruses; however, *Ae. albopictus* was largely responsible for the dengue and chikungunya outbreaks in Gabon in 2007 and 2010 [14, 17, 21]. Furthermore, *Ae. albopictus* is able to transmit the chikungunya virus variant possessing the E1-226V mutation more efficiently than *Ae. aegypti* [22, 23]. This mutation was first identified during the chikungunya outbreak in the African Indian Ocean islands in 2005 [24], and was later isolated in central Africa [18, 19, 25].

In DRC, 50,000 suspected cases were reported during the first chikungunya outbreaks in Kinshasa from 1999 to 2000 [16]. Chikungunya outbreaks also occurred in Kinshasa in 2012 and 2019 and in the adjacent Kongo Central Province in 2019 [25, 26]. In addition, the number of dengue virus infections has also increased in recent years [26-29]. Although an apparent outbreak did not occur, an entomological study caught several *Aedes* mosquitoes infected with CHIKV in Kinshasa in 2014 [30]. Moreover, a study confirmed involvement of *Ae. albopictus* for transmitting CHIKV with the E1-A226V mutation in two cities, Matadi and Kasangulu, of Kongo Central Province during the 2019 chikungunya outbreak [25].

Curative treatments and vaccines are not available for dengue and chikungunya [31, 32], and thus vector control is a valuable available tool for reducing infections [ 33]. As such, understanding the current distribution of *Ae. albopictus* in DRC is an essential step for the control. Global level distribution models based on environmental variables indicate that almost the entire area of DRC is suitable for *A. albopictus* establishment [3, 34, 35]. These models were constructed without entomological data from DRC, and thus the provided information was too coarse to apply to local vector control. In the present study, we described the current distribution of *Ae. albopictus* in DRC based on locally available data. In particular, we provided detailed information for Kinshasa and the neighboring areas where chikungunya outbreaks recently occurred. We also revealed important environmental variables related to the distribution, and attempted to determine if the present distribution is static.

## Materials and methods

### Study areas

DRC is the largest country in Sub-Saharan Africa with an area of roughly 2,4 million km^2^, and possesses a diversity of landscapes and climates. The country is divided into six geographic regions (western, northern, far-northern, central, eastern, and southeastern) based on landscape and climate (Fig 1). The landscape of the western region is composed of the coastal plain, with hills and plateaus in the south. The vegetation type is mainly savannah, with a tropical humid climate and a three 3-month dry season. This region includes Kinshasa and Kongo Central province, where chikungunya and dengue outbreaks have occurred. The Congo Basin and equatorial forests largely occupy the northern region. This region has an equatorial climate without a dry season. The far-northern region is characterized with savannahs, and has a tropical humid climate with a three month dry season. Equatorial forests occupy the northern part of the central region, whereas the southern part is mainly plateau with savannahs and steppes. The central region has a dry tropical climate with a three month dry season. High hills and mountains dominate the eastern region, and lush vegetation forms the mountain forests. The region has a temperate mountain climate without a distinct dry season. The southeastern region is dominated by high plateaus with savannahs. The region has a dry tropical climate with a six-month dry season.

**Fig 1.**
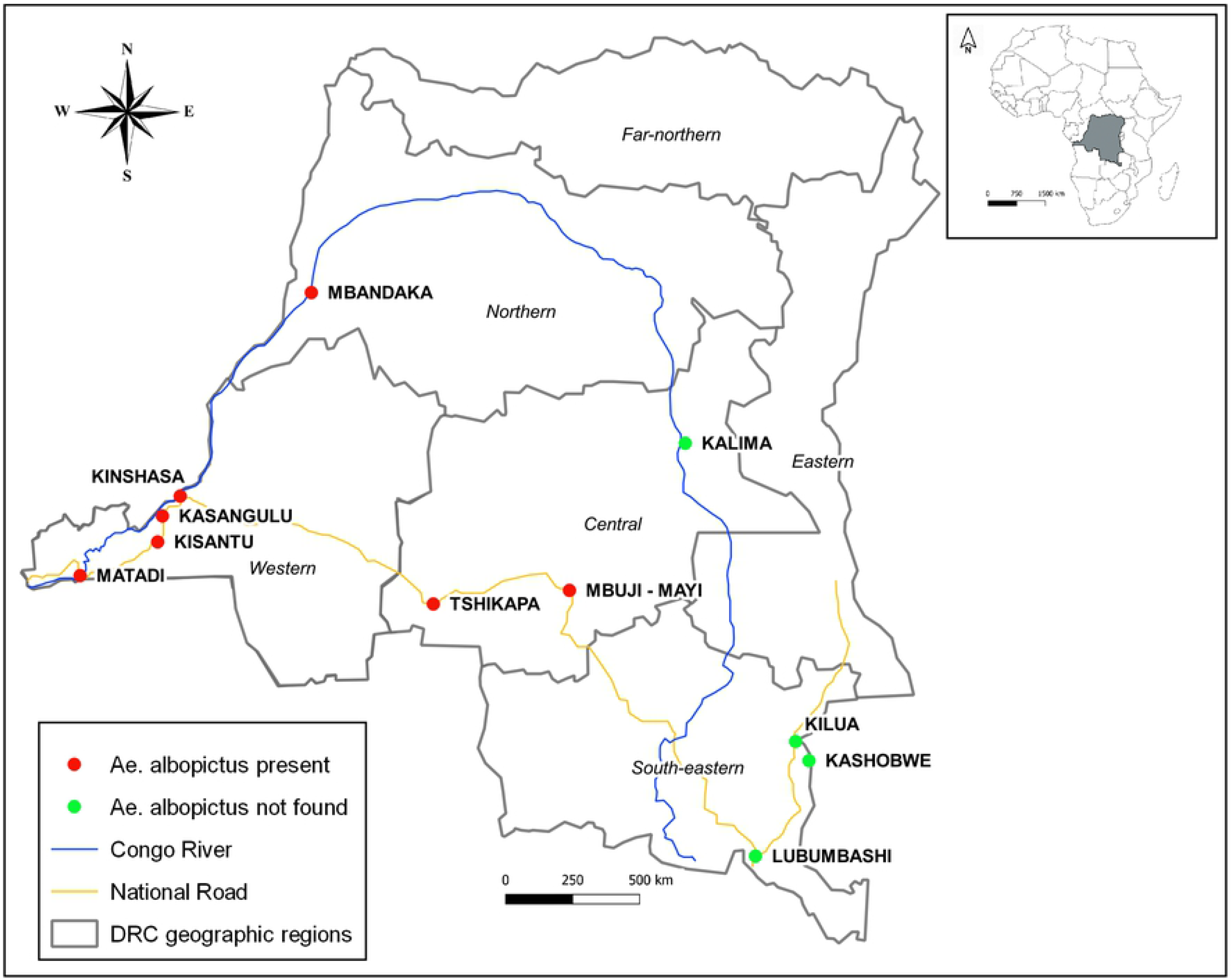
Distribution of *Ae. albopictus* in DRC. Red dots depict the presence of *Ae. albopictus*, and green dots depict absence at the city level. Mosquitoes were sampled at several sites within Matadi, Kisantu, Kasangulu, Kinshasa, and Mbandaka, and *Ae. albopictus* was found at one site at least. Each geographic region is made up of multiple provinces, represented by boundaries.

We conducted entomological surveys at 32 sites within 11 cities across four different geographic regions except the eastern and far-northern regions, from May 2017 to September 2019 (Table 1). First, we focused on the western region in which *Ae. albopictus* has been recorded [13, 25]. The survey in the western region included 14 sites within Kinshasa and nine sites in the three cities, Kasangulu, Kisantu, and Matadi, in Kongo Central Province. Since human-mediated dispersal of *Ae. albopictus* was an immediate concern, the survey also included nine sites along the major transportation routes (Congo River and national roads) in the other three regions (Fig 1). These sites were three sites within Mbandaka in the western part of the northern region; Tshikapa, Mbuji-Mayi, and Kalima in the central region and Lubumbashi, Kilwa, and Kashobwe in the southeastern region.

**Table 1.**
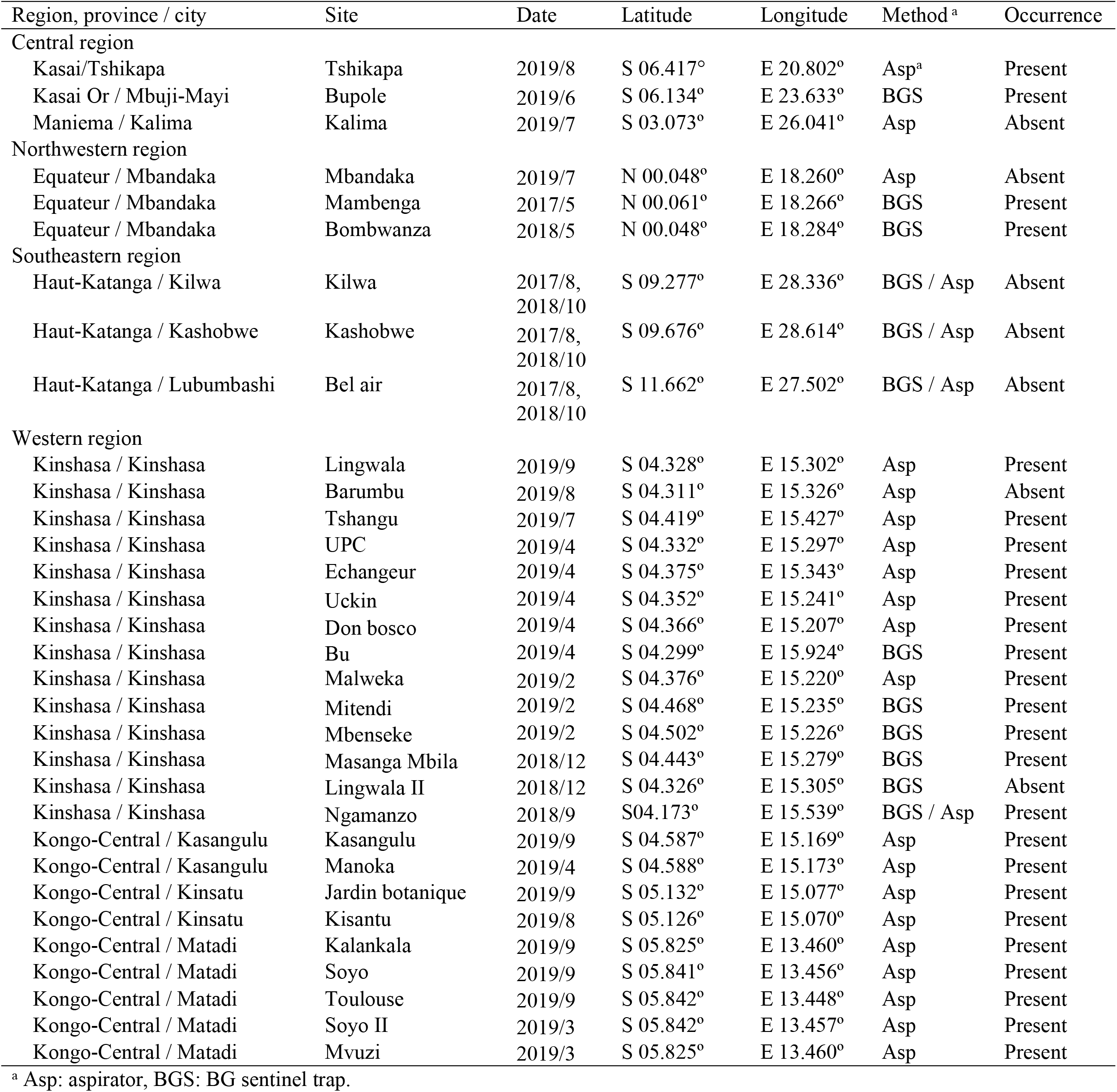
Sampling sites, methods and occurrence of *Ae. albopictus*.

| Region, province / city | Site | Date | Latitude | Longitude | Method <sup>a</sup> | Occurrence |
| --- | --- | --- | --- | --- | --- | --- |
| Central region |  |  |  |  |  |  |
| Kasai/Tshikapa | Tshikapa | 2019/8 | S 06.417° | E 20.802° | Asp<sup>a</sup> | Present |
| Kasai Or / Mbuji-Mayi | Bupole | 2019/6 | S 06.134° | E 23.633° | BGS | Present |
| Maniema / Kalima | Kalima | 2019/7 | S 03.073° | E 26.041° | Asp | Absent |
| Northwestern region |  |  |  |  |  |  |
| Equateur / Mbandaka | Mbandaka | 2019/7 | N 00.048° | E 18.260° | Asp | Absent |
| Equateur / Mbandaka | Mambenga | 2017/5 | N 00.061° | E 18.266° | BGS | Present |
| Equateur / Mbandaka | Bombwanza | 2018/5 | N 00.048° | E 18.284° | BGS | Present |
| Southeastern region |  |  |  |  |  |  |
| Haut-Katanga / Kilwa | Kilwa | 2017/8,<br>2018/10 | S 09.277° | E 28.336° | BGS / Asp | Absent |
| Haut-Katanga / Kashobwe | Kashobwe | 2017/8,<br>2018/10 | S 09.676° | E 28.614° | BGS / Asp | Absent |
| Haut-Katanga / Lubumbashi | Bel air | 2017/8,<br>2018/10 | S 11.662° | E 27.502° | BGS / Asp | Absent |
| Western region |  |  |  |  |  |  |
| Kinshasa / Kinshasa | Lingwala | 2019/9 | S 04.328° | E 15.302° | Asp | Present |
| Kinshasa / Kinshasa | Barumbu | 2019/8 | S 04.311° | E 15.326° | Asp | Absent |
| Kinshasa / Kinshasa | Tshangu | 2019/7 | S 04.419° | E 15.427° | Asp | Present |
| Kinshasa / Kinshasa | UPC | 2019/4 | S 04.332° | E 15.297° | Asp | Present |
| Kinshasa / Kinshasa | Echangeur | 2019/4 | S 04.375° | E 15.343° | Asp | Present |
| Kinshasa / Kinshasa | Uckin | 2019/4 | S 04.352° | E 15.241° | Asp | Present |
| Kinshasa / Kinshasa | Don bosco | 2019/4 | S 04.366° | E 15.207° | Asp | Present |
| Kinshasa / Kinshasa | Bu | 2019/4 | S 04.299° | E 15.924° | BGS | Present |
| Kinshasa / Kinshasa | Malweka | 2019/2 | S 04.376° | E 15.220° | Asp | Present |
| Kinshasa / Kinshasa | Mitendi | 2019/2 | S 04.468° | E 15.235° | BGS | Present |
| Kinshasa / Kinshasa | Mbenseke | 2019/2 | S 04.502° | E 15.226° | BGS | Present |
| Kinshasa / Kinshasa | Masanga Mbila | 2018/12 | S 04.443° | E 15.279° | BGS | Present |
| Kinshasa / Kinshasa | Lingwala II | 2018/12 | S 04.326° | E 15.305° | BGS | Absent |
| Kinshasa / Kinshasa | Ngamanzo | 2018/9 | S04.173° | E 15.539° | BGS / Asp | Present |
| Kongo-Central / Kasangulu | Kasangulu | 2019/9 | S 04.587° | E 15.169° | Asp | Present |
| Kongo-Central / Kasangulu | Manoka | 2019/4 | S 04.588° | E 15.173° | Asp | Present |
| Kongo-Central / Kinsatu | Jardin botanique | 2019/9 | S 05.132° | E 15.077° | Asp | Present |
| Kongo-Central / Kinsatu | Kisantu | 2019/8 | S 05.126° | E 15.070° | Asp | Present |
| Kongo-Central / Matadi | Kalankala | 2019/9 | S 05.825° | E 13.460° | Asp | Present |
| Kongo-Central / Matadi | Soyo | 2019/9 | S 05.841° | E 13.456° | Asp | Present |
| Kongo-Central / Matadi | Toulouse | 2019/9 | S 05.842° | E 13.448° | Asp | Present |
| Kongo-Central / Matadi | Soyo II | 2019/3 | S 05.842° | E 13.457° | Asp | Present |
| Kongo-Central / Matadi | Mvuizi | 2019/3 | S 05.825° | E 13.460° | Asp | Present |

<sup>a</sup> Asp: aspirator, BGS: BG sentinel trap.

### Mosquito sampling

Within each site, sampling was focused on places around dwellings which are ecologically suitable for adults of *Ae. albopictus*, and places where residents reportedly experience frequent day-time mosquito bites. *Aedes* mosquitoes were collected with electric aspirators (Prokopack Aspirator, John W. Hock, Gainesville, USA) and/or BG sentinel traps (Biogents Inc, Regensburg, Germany) from 3:00 pm to 6:00 pm for three to seven consecutive days at each site. Sampled mosquitoes were identified morphologically to species according to Huang’s identification keys [36]. When at least one *Ae. albopictus* was collected, the site was considered as a positive site. A distribution map was constructed using the Quantum Geographic Information System software version 3.4.13 (QGIS Development Team, 2020) (Fig 1).

### Environmental variables

We reviewed literature related to modelling *Ae. albopictus* distribution using the maximum entropy software, MaxEnt [37]. This software is often used for modeling species distribution, and effectively handles a small number of collection sites [38-42]. Based on the review, we selected 18 environmental variables which had a permutation importance (PI) of at least 5% (Table 2) [34, 43-55]. PI indicates the importance of each variable in a MaxEnt model [56]. Among the 18 variables, 15 climatic variables were obtained from the WorldClim database (http://www.worldclim.com/version2) [57]. This climate database provides average historical climate data from 1970 to 2000 with a spatial resolution of 1 km x 1 km. Digital elevation model (DEM) data was obtained from SRTM imagery/USGS with a resolution of 30.9 m (or 1-arc second) (https://www2.jpl.nasa.gov/srtm/). The datasets of two vegetation variables, Enhanced Vegetation Index (EVI) and Normalized Differentiation Vegetation Index (NDVI), were downloaded from Modis Vegetation Index/USGS with a resolution of 1km x 1 km (https://modis.gsfc.nasa.gov/data/dataprod/mod13.php). Dry season length was included in addition to the variables obtained by the literature review [58].

**Table 2.** Important environmental variables for *Aedes albopictus* distribution.

| Code | Variable | PI (%) | References |
| --- | --- | --- | --- |
| Bio1 | Annual mean temperature |  | [34, 47, 54] |
| Bio2 | Mean diurnal temperature range | 55.6 | [45, 49] |
| Bio4 | Temperature seasonality |  | [43, 49] |
| Bio5 | Maximum temperature of warmest month | 30.8 | [43, 45, 47] |
| Bio6 | Minimum temperature of coldest month |  | [43, 47] |
| Bio7 | Temperature annual range |  | [45] |
| Bio10 | Mean temperature of warmest quarter |  | [48, 50, 52, 53] |
| Bio11 | Mean temperature of coldest quarter |  | [46, 48, 50, 52, 53, 55] |
| Bio12 | Annual precipitation |  | [47, 55] |
| Bio13 | Precipitation of wettest month |  | [43, 45, 47, 51] |
| Bio14 | Precipitation of driest month |  | [44, 47, 49] |
| Bio15 | Precipitation seasonality |  | [43] |
| Bio16 | Precipitation of wettest quarter |  | [46] |
| Bio17 | Precipitation of driest quarter |  | [46, 50, 55] |
| Bio18 | Precipitation of warmest quarter |  | [49, 54] |
| DEM | Digital elevation model |  | [54] |
| NVDI | Normalized Difference Vegetation Index |  | [34] |
| EVI | Enhanced vegetation index | 13.6 | [55] |
|  | Dry season length |  | [58] |

Permutation importance values are given for variables selected in the final model.

### Modeling

We selected environmental variables that were significantly different between positive and negative sites. A relationship of mosquito occurrence with each variable was examined using the Wilcoxon-Mann-Whitney test (GraphPad Prism version 8.4.2, GraphPad Software, San Diego, California USA). When numbers of sample size were insufficient (n < 4) for the statistical test, we identified variables which had an extreme median value at negative sites versus positive sites. We first examined if a negative site median value was within the range of positive site values in the corresponding geographic region. When the median value was outside the range, we also compared it to the range of values from all positive sites including ones from the other regions. When the value was still outside the range, the variable was considered for modeling.

Between the selected variables, we examined the Pearson correlation coefficients [44]. When the coefficients were above 80%, we retained them based on their apparent importance in past studies (Table 2) [34, 43-55]. Dry season length was excluded from the analyses because of the absence of a raster file. Then, we ran a full model including all selected variables with the default settings of MaxEnt. Based on the results from the full model, we constructed a reduced model including variables that had a PI above 5%. Since our sample size was small, we modified the settings in MaxEnt using ten replications, linear feature, and cumulative output format. The PI from the latter model was used to identify the most important variables. Response curves were also used to determine how the model changes with a permutation of each variable. The area under the curve (AUC) was used to assess model accuracy. When an AUC value was above 0.75, the model was acceptable. With the outputs from the optimal model, we constructed a predicted geographical distribution map of *Ae. albopictus* in DRC using the QGIS software.

## Results

We collected a total of 2,841 *Aedes* mosquitoes. Of which, 2,331 (82%) were *Ae. albopictu*s, and 510 (18%) were *Ae. aegypti*. The former specie*s* was found at 25 of 32 sites within 7 of 11 cities (Table 1, Fig 1). Within Kinshasa, *Ae. albopictus* was collected at 12 of 14 sites (Table 1). In Kongo Central Province, *Ae. albopictus* was collected at all nine sites. This species was collected at two of the three sites within one city in the western part of the northern region. In the central region, we found *Ae. albopictus* in the two cities in the southern part, Tshikapa and Mbuji-Mayi, but we did not find it in the city in the northeastern part, Kalima. We did not find *Ae. albopictus* in the three cities, Kilwa. Kashobwe and Lubumbashi, in the southeastern region (Table 1).

A total of 19 environmental variables were selected based on a literature review (Table 2). Wilcoxon-Mann-Whitney tests revealed that the precipitation of the warmest quarter was significantly greater at the positive sites compared with the negative sites; however, the differences were not statistically significant for the other variables (Fig 2). The medians of all environmental variables at the two negative sites in the western region were within the ranges of values at the positive sites of the same region (Fig 3). In the northern region, the medians from the negative sites were within the range of values from the positive sites except for the NVDI (Fig 3R). However, the median of NDVI was within the range of the values from the positive sites when all regions were considered. The medians of nine variables at the negative site in the central region were out of the ranges of the two positive sites. When all regions were considered, the medians were within the range of the positive sites. However, the maximum temperature of the warmest month at the negative site in the central region was lower than the range of all positive site values including ones from the other regions. The same negative site of the central region had higher EVI and NDIV than the ranges of all positive sites. The medians of ten variables at the three negative sites in the southeastern region were outside the ranges of values at the positive sites. The negative sites had lower annual mean temperatures, a wider mean diurnal temperature range, lower minimum temperatures of the coldest month, lower mean temperatures of the coldest quarter, a wider temperature annual range, greater precipitation seasonality, lower precipitation of the driest quarter, lower precipitation of the warmest quarter, higher elevation, and longer dry season length than any of the positive sites. Lubumbashi is located in the southernmost and at the highest elevation among the sites in the southeastern region, and these environmental variables of the city were more extreme than the other sites.

**Fig 2.**
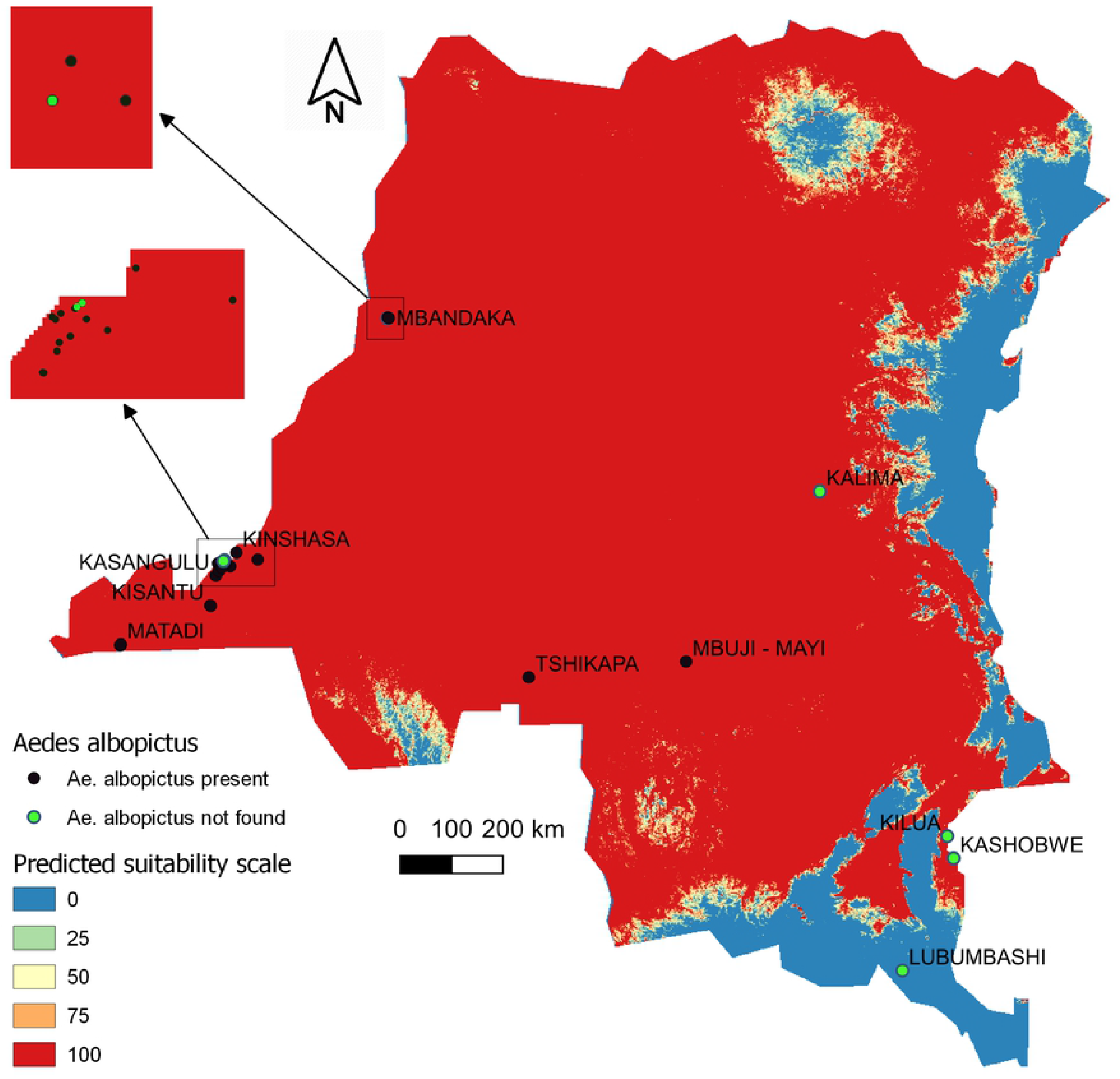
Comparisons of each environmental variable between the positive and negative *Ae. albopictus* collection sites. Each panel shows the first quartile, the median, the third quartile, the minimum and the maximum values in positive (*Ae. albopictus* was found) and negative (the species was not found) sites by box plots. A: Annual mean temperature (°C); B: mean diurnal temperature range (°C); C: temperature seasonality (%); D: maximum temperature of warmest month (°C); E: minimum temperature of the coldest month (°C); F: mean temperature of the coldest quarter (°C); G: temperature annual range (°C); H: mean temperature of the warmest quarter (°C); I: annual precipitation (mm); J: precipitation of the wettest month (mm); K: precipitation of the driest month (mm); L: precipitation seasonality (%); M: precipitation of the wettest quarter (mm); N: precipitation of the driest quarter (mm); O: precipitation of the warmest quarter (mm); P: digital elevation model (m); Q: enhanced vegetation index; R: normalized difference vegetation index; S: dry season length (month). An asterisk indicates that that the difference was statistically significant (p < 0.05) with Wilcoxon-Mann-Whitney tests.

**Fig 3.**
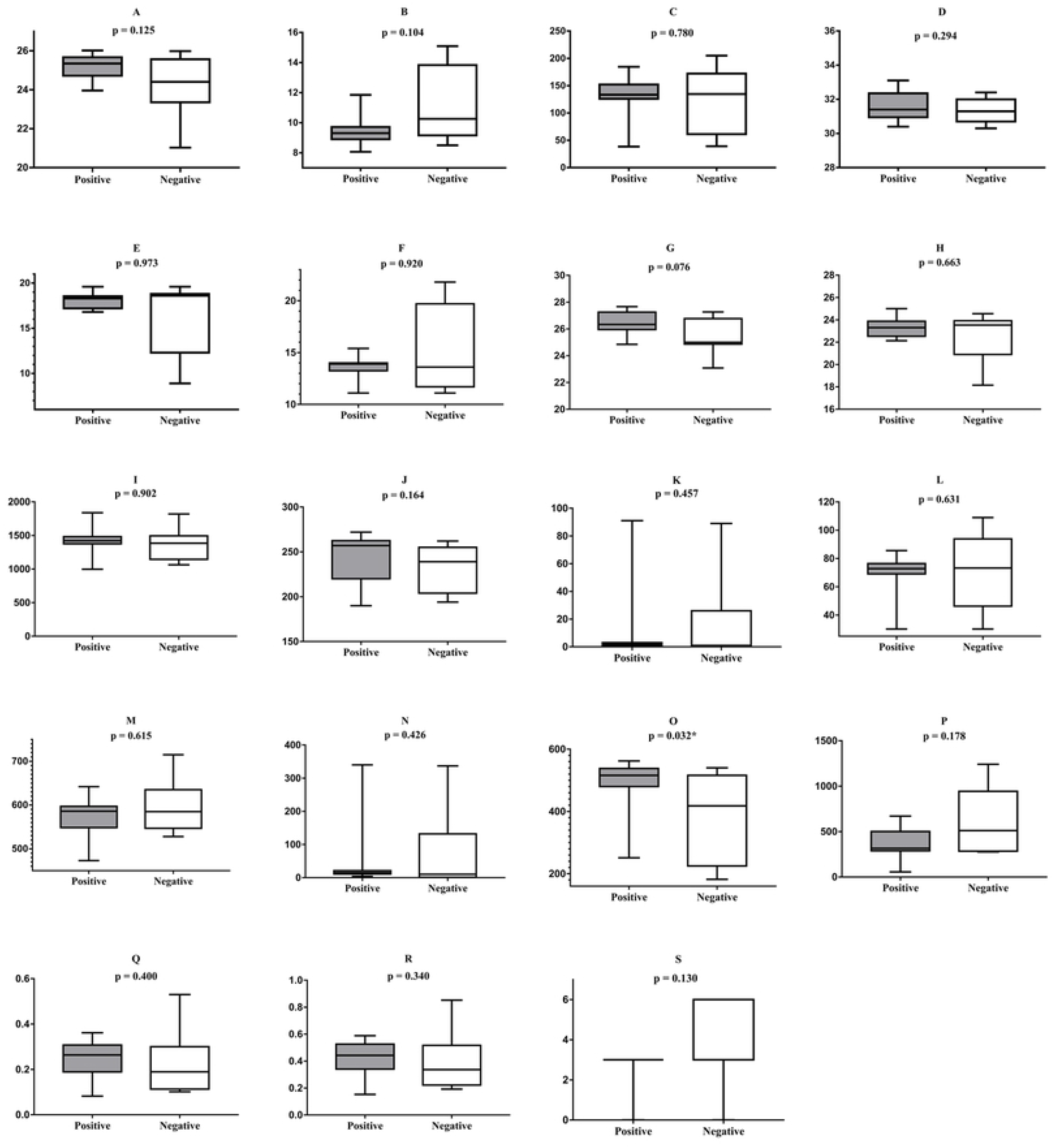
Medians of each environmental variable at positive sites and negative sites of *Ae. albopictus* in the four regions. A value for each site is depicted as a dot. The black horizontal bars indicate the median and vertical bars indicate the range. A: annual mean temperature (°C); B: mean diurnal temperature range (°C); C: temperature seasonality (%); D: maximum temperature of warmest month (°C); E: minimum temperature of the coldest month (°C); F: mean temperature of the coldest quarter (°C); G: temperature annual range (°C); H: mean temperature of the warmest quarter (°C); I: annual precipitation (mm); J: precipitation of wettest month (mm); K: precipitation of the driest month (mm); L: precipitation seasonality (%); M: precipitation of the wettest quarter (mm); N: precipitation of the driest quarter (mm); O: precipitation of the warmest quarter (mm); P: digital elevation model (m); Q: enhanced vegetation index; R: normalized difference vegetation index; S: dry season length (month).

Of 12 selected variables, excluding dry season length, five pairs were highly correlated among eight variables (S1 File). We chose annual mean temperature, mean diurnal temperature range and the EVI over the others because the past studies showed that they were more important. As a result, seven variables were included in the full MaxEnt analysis (Table 2). After the model selection, the optimal model contained three variables, maximum temperature of the warmest month, mean diurnal temperature range, and EVI. Mean diurnal temperature range was the most important variable, followed by maximum temperature of warmest month, and EVI (Table 2). The AUC of the optimal model was 0.975. The response curves revealed that the highest suitable area was predicted with EVI below – 0.017, maximum temperature of the warmest month above 34.3^°^C, and mean diurnal temperature below 6.5°C (Fig 4).

**Fig 4.**
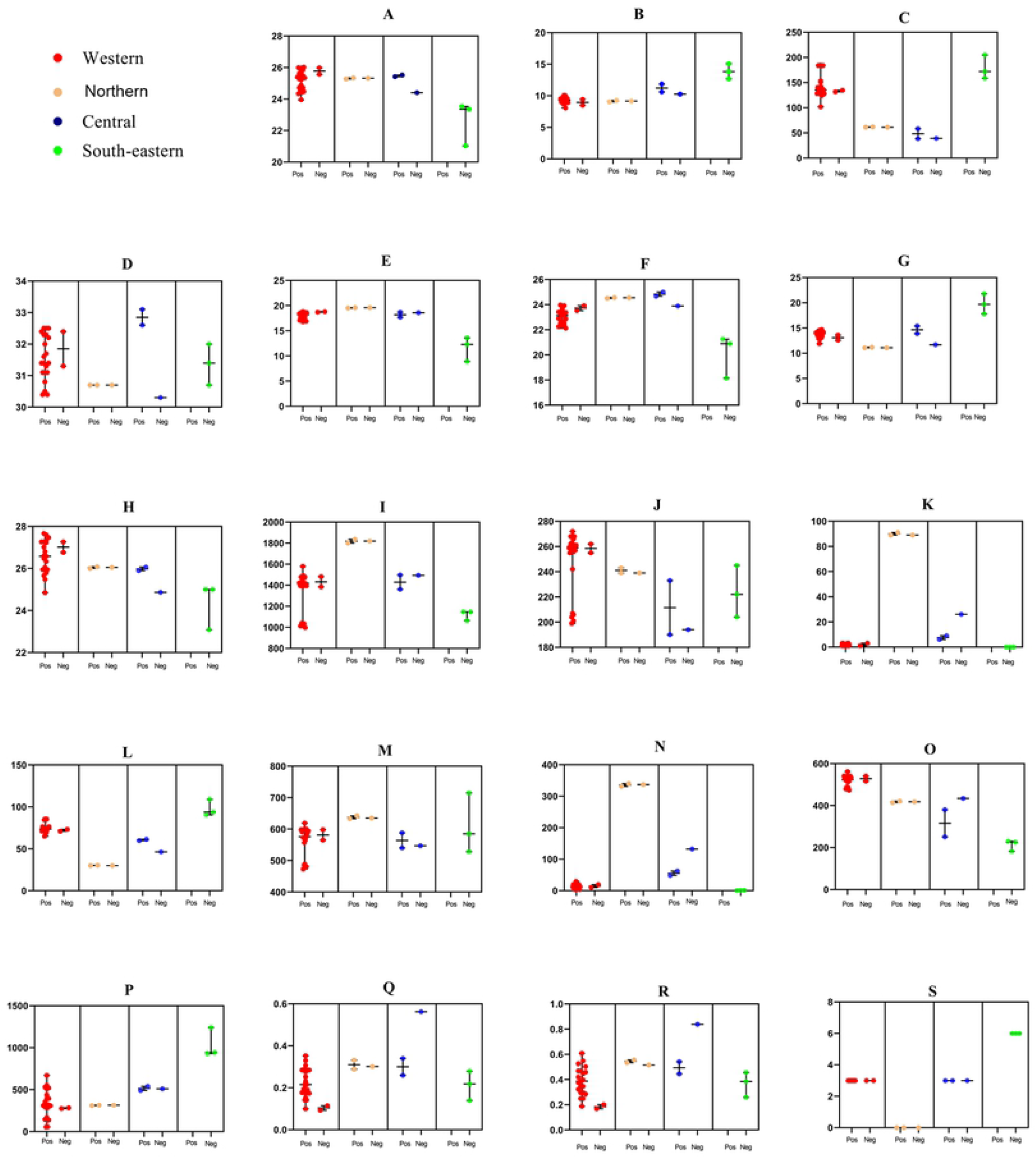
Response curves for *Ae. albopictus* suitability in relation to mean diurnal temperature range (A), maximum temperature of warmest month (B), and enhanced vegetation index (C). The curves show how each environmental variable affects the MaxEnt prediction. The red line is the mean response of the ten MaxEnt replications.

The model predicted that most of DRC is suitable for *Ae. albopictus* establishment (Fig 5). The suitability was high in the most parts of the western region; however, it varied between 0 to 75% in the southern area of the region. The suitability was also high in the central region and the northern region although a noticeable area in the northeastern region had low suitability. The eastern part of the eastern region and the southern part of the southeastern region had low suitability. The model successfully predicted all positive sites within the highly suitable areas and all negative sites within the highly suitable areas in the western, the northwestern, and the central regions. However, the model predicted two negative sites, Kilwa and Kashobwe, in the southeastern region to be suitable whereas Lubumbashi was predicted as being unsuitable area.

**Fig 5.**
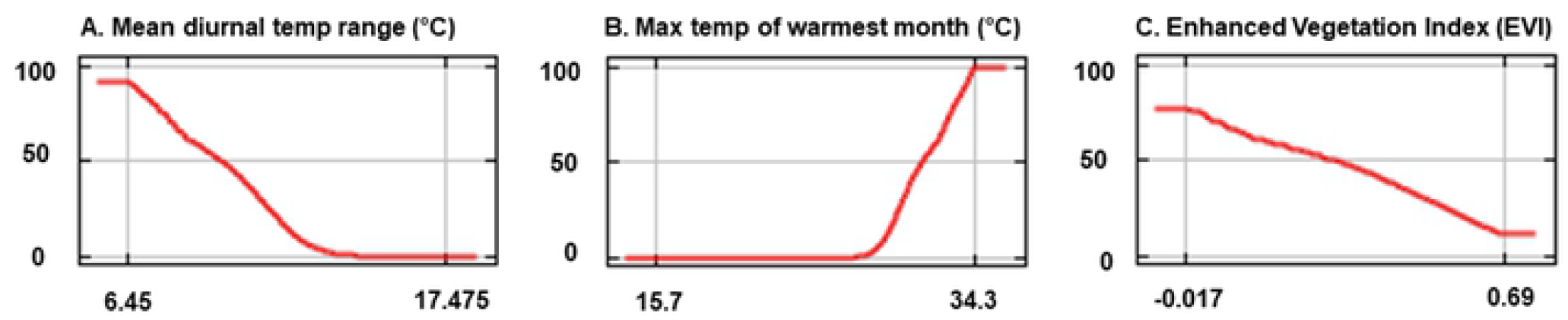
Suitability map of *Ae. albopictus* in DRC generated by the optimal MaxEnt model. Dots depict the presence (black) or absence (green) of *Ae. albopictus*. Only 24 out of the 32 dots can be visualized because some sites are overlapped.

## Discussion

The present study found *Aedes albopictus* in 25 sites in seven cities in DRC. This mosquito species was newly found in four cities in the western and central regions, but it was absent in the cities in the southeastern region where many environmental variables showed extreme values. The MaxEnt model revealed that the occurrence of *Ae. albopictus* was positively associated with maximum temperature of the warmest month, and negatively with wider mean diurnal temperature range and enhanced vegetation index. The model predicted that almost the entire area of DRC is suitable for the establishment of *Ae. albopictus*.

Within Kinshasa, *Ae. albopictus* was found at 12 of 14 collection sites. This mosquito species was recorded in Kinshasa for the first time in DRC in 2016 [13]. A recent study reported within this city a high level of larval infestation of *Ae. albopictus* in artificial containers together with *Ae. aegypti* [59]. In the adjacent province, *Ae. albopictus* was found at all 9 collection sites within three cities, Kasangulu, Kisantu, and Matadi. The present study recorded this mosquito species in Kisantu for the first time, while it was recorded in Matadi and Kasangulu during the 2019 chikungunya outbreak. During the outbreak, *Ae. albopictus* was more abundant than *Ae. aegypti* in these two cities [25]. The findings from the present study were sufficient to conclude that *Ae. albopictus* is well established in the western part of the western region.

We also confirmed that *Ae. albopictus* has extended its distribution to the inland cities. This mosquito species was recorded in Mbandaka in the northern region for the first time. We collected *Ae. albopictus* in the city in 2017 and in the two consecutive years, indicating that this mosquito quickly spread to the area after its recording in Kinshasa in 2016. This species was likely introduced to Mbandaka from the western region by traffic along the Congo River, which is the main transportation route to the northern region. In the Philippines a molecular study showed evidence of *Ae. aegypti* migrations with ships among the islands [60].

In contrast, we did not find *Ae. albopictus* in Kalima in the upriver region of the Congo River in the eastern part of the central region. The result is likely due to the distance and the poor access from the other areas where this species has become established. However, air flight activity is intense between the area and Kinshasa, and *Ae. albopictus* might be introduced by air in the future [61]. Either way, the result from one collection site is not enough to confirm the absence of this mosquito species in the region. On the other hand, *Ae. albopictus* was found at two cities in the southern part of the central region. The results are likely due to a larger amount of traffic and a shorter distance between Kinshasa and this area compared with Kalima. The access is also better through the major roads, and there are frequent flights between Kinshasa and the area.

We did not find *Ae. albopictus* at all three cities in the southeastern part of the southeastern region. The results may be partially due to the distances from the areas where this mosquito has been established. However, because Lubumbashi is the second largest city in DRC, the amount of road traffic from the central and western regions is not negligible, and the flight activities are intense between Kinshasa and Lubumbashi. The intense traffic may introduce this mosquito species to the area in the near future [62].

Climate may limit the distribution of *Ae. albopictus* in the southeastern region. The medians of ten environmental variables at the negative sites in the southeastern region were outside the ranges of the values from the positive sites of the other regions. The results indicate that the sites in the southeastern region are cooler, and the temperature fluctuates more because of the inland with high altitudes. Indeed, the MaxEnt model indicated that the climate variables (maximum temperature of the warmest month and mean diurnal temperature range) are important for establishment of this mosquito species. On the other hand, the model suggests that two negative sites, Kilwa and Kashobwe, in the southeastern region are suitable for *Ae. albopictus* establishment. The elevations of these sites are less than 1,000 m, the maximum temperature of warmest months is 31 to 32 °C and the mean annual temperatures are 23 to 24 °C. Since *Ae. albopictus* could establish in temperate areas with an annual mean temperature of 11°C and/or 1,350 accumulated degree-days above 11°C per year [63-65], the temperatures of the two cities are warm enough. These model results suggest that the distances and traffic from the western region are likely the limiting factors, but this mosquito species may establish in these two sites in the future.

The model suggests that Lubumbashi is not suitable for *Ae. albopictus* survival. This city is situated at an elevation of about 1,200 m, and the mean annual temperature is 21°C. While the maximum temperature of the warmest month is 31°C, the minimum temperature of the coldest month, July, drops to 9 °C. The coldest month occurs in the middle of the six-month dry season when the monthly rainfall often becomes less than 1 mm. While the lengths of the dry season are similar among the three cities in the region, the lower temperature and wider diurnal temperature range may make the climate condition of Lubumbashi less favorable for *Ae. albopictus*. Even though eggs of this mosquito are tolerant to desiccation [66], egg survivorship would become less with decreases of temperature and humidity during the dry season [60, 67]. Furthermore, a greater fluctuation of temperature may make the conditions less favorable for survival [67, 68]. The conditions may become even tougher for *Ae. albopictus* strains originating from tropic regions, which are less tolerant to cooler climate compared with strains from temperate regions [69, 70].

A study in Madagascar reported that the distribution of *Ae. albopictus* is largely limited to the eastern part of the island, with high humidity, a temperature of the coldest months above 12 °C, and dry season shorter than six months in length [58]. The study, however, found *Ae. albopictus* breeding in used tires and captured adults in residential areas in the southwestern region with an annual precipitation less than 600 mm and an eight-month dry season. The findings in Madagascar suggest that this mosquito species is able to establish in an area where suitable man-made habitats are available as long as the temperature is warm enough. Although *Ae. albopictus* distribution in Asia, from which it originated, occurs more in rural areas with greater vegetation, it also utilizes artificial habitats such as discarded containers in urban areas [1, 13]. Probably the entry point of a new region is likely an urban area with a larger amount of traffic. This partially explains the negative association of this species with the enhanced vegetation index indicated by the MaxEnt model.

Although our field survey did not cover the far-northern region and the eastern region, the model suggests that most of the far-northern region and the western part of the eastern region are also suitable for establishment of this mosquito species. *Ae. albopictus* might have already reached these regions, or it may reach there in the near future. In contrast, the model suggests that the eastern part of the eastern region is not suitable for this mosquito species. The area is 2,000 m above sea level, and includes mountains above 4,000 m. The harsh climate likely does not allow *Ae. albopictus* to establish in the area [63-65].

## Limitation

The number of collection sites was small relative to the size of the country. Including the far-northern region and the eastern region, a larger number of collection sites could provide a better picture of the relationships of *Ae. albopictus* with the environmental variables. The Wilcoxon-Mann-Whitney tests revealed that precipitation of the warmest quarter was greater at the positive sites than the negative sites. Although this is the only variable statistically different between them, other variables might become significant with a larger number of collection sites.

We collected mosquitoes mainly within urban areas. Mosquitoes are more frequently introduced to urban areas with human activities, and thus sampling approach was practical to identify sites in which *Ae. albopictus* was established when considering the large size of the country. For instance, with fewer negative sites, the Max Ent model might be affected by the highest EVI value at the single negative site in the central region. As a result, EVI became one of the three important environmental variables, and it was negatively associated with the presence of *Ae. albopictus*. This result contradicts the past studies in the other areas [71]. A more precise picture would be produced with a finer spatial scale which can recognize small patches of vegetation within an urban area, though it is still challenging with free satellite data.

The environmental variables used in the present study were selected based on studies conducted mostly in temperate areas, because few studies were conducted in Africa. Appropriate variables for the African situation might be different.

## Conclusion

*Aedes albopictus* has established populations in the major cities of the western region of DRC. This mosquito species is expanding its geographical distribution toward the inland. The migration is likely facilitated by the major transportation routes including the Congo River. The MaxEnt model based on environmental variables suggests that most of the country is suitable for the establishment of *Ae. albopictus*, except the areas in the eastern and the southeastern parts of the country. The results from our study suggest that low temperatures and a long dry season limit the distribution of *Ae. albopictus*. This is the first report to provide the current and future *Ae. albopictus* distributions in DRC using locally collected mosquito data.

## Implication

Autochthonous cases of chikungunya and dengue have been reported from the western region and the southern part of the central region where we found *Ae. albopictus* [72]. Although *Ae. albopictus* was found in the southwestern part of the northern region, autochthonous cases of the viral diseases have not been reported. The diseases have not been reported from the northern part of the central region and the southeastern region where we did not find this mosquito species. Moreover, the diseases have not been reported from the far-northern area and the eastern region. Our model implies that, following the expansion of mosquito distribution, chikungunya and dengue may also spread to most parts of the country in the near future. Country-wide entomological surveillance is needed to detect the signs of impending epidemics.

## Acknowledgments

We would like to thank Pitshou Mampuya, Herve Michel Vulu and Kelvin Landu for their helpful contribution in the field activities. Fabien Vulu is grateful to the Program for Nurturing Global Leaders in Tropical and Emerging Communicable Diseases, Graduate school of Biomedical Sciences, Nagasaki University, and Japan International Cooperation Agency (JICA) for their support.

## Supporting file captions

**S1 File. Pearson correlations between environmental variables**.

**S1 Dataset. Environmental variable data from all collection sites**.

